# An exact formula for the contribution of sampling error to *r*^2^, a common measure of linkage disequilibrium

**DOI:** 10.64898/2026.05.19.726388

**Authors:** Robin S. Waples

**Author notes:** Corresponding author: Robin Waples.

## Abstract

Interest in quantifying linkage disequilibrium (LD, non-random associations of alleles at different loci) has skyrocketed in recent years as researchers have focused on use of LD in genome-wide association studies (GWAS), for studying historical demography, and for estimating effective population size (*N*_*e*_). The most widely used LD metric is *r*_2_ = the squared correlation of alleles at a pair of loci. Despite a half century of efforts, developing an unbiased expectation of *r*^2^ as a function of the many factors that can affect it (physical linkage, genetic drift, selection, migration, mutation, mating systems) remains elusive. Furthermore, even when all of these other factors are absent, empirical estimates of *r*^2^ are upwardly biased by sampling a finite number (*S*) of individuals, and that must be accounted for if one wants to focus on the desired signal of LD. Previous approaches to estimate 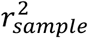 have been shown to be biased to greater or lesser degrees. The purpose of this short paper is to demonstrate that a simple and apparently exact expression for 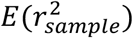 does exist for the special case where sampling error is the only factor contributing to *r*^2^, in which case 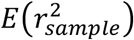 = 1/(*S* − 1). When other factors contribute heavily to LD, 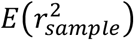 shrinks toward 0 as empirical *r*^2^ → 1. However, for estimating contemporary *N*_*e*_ with unlinked markers, empirical *r*^2^ will generally be small and 1/(*S* − 1) will provide a robust estimate of 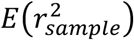.

## Introduction

Linkage disequilibrium (LD, non-random associations of alleles at different gene loci) has attracted the attention of evolutionary biologists for over a century (Slatkin 2008; Sved and Hill 2018). Interest in LD has intensified in recent decades for several major reasons: tightly linked loci hold considerable promise for identifying genes associated with human diseases (Bulik-Sullivan et al. 2015a); linked loci also provide a window into the past by providing insights into historical demography (Patterson et al. 2012; Santiago et al. 2020); LD between loci on different chromosomes is now commonly used to estimate contemporary effective population size (*N*_*e*_) in natural populations of diverse taxa (Waples and Do 2008; Clarke et al. 2024).

A widely-used approach for quantifying LD is to calculate the squared correlation coefficient between alleles at two loci, *r*^2^. Over half a century of efforts to develop a theoretical expectation for *r*^2^ have met with limited success (Ohta and Kimura 1969; Sved and Feldman 1973; Weir and Hill 1980; Golding 1984; Hudson 1985; Sved et al. 2013). Three related issues make this a hard problem. First, even if we restrict consideration to selectively neutral genes in closed, randomly-mating populations, several different factors (and their interactions) can contribute to observed levels of LD: reproduction by a finite effective number of parents 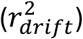, lack of independence of syntenic genes 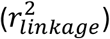, and sampling error 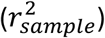. Second, conceptually, *r*^2^ for a population of interest applies to an infinite array of progeny produced by the parents (Weir and Hill 1980; hereafter W&H). In the real world, of course, all samples of individuals are finite, and often small in number, as researchers increasingly focus on assaying larger numbers of genes in fewer individuals. Finite samples create an upward bias in *r*^2^ that must be accounted for if one wants to focus on the desired signal of LD. Finally, *r*^2^ is ratio, so it is likely that no completely unbiased expectation exists (Bercovich et al. 2026). The typical approach is to approximate *E*(*r*^2^) as a ratio of expectations. The inability to derive a completely unbiased expectation for the contribution of sampling error to 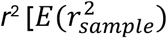 limits the accuracy of inferences regarding underlying evolutionary processes.

The purpose of this short paper is to demonstrate that a simple and apparently exact expression for 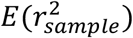 does exist for the special case where sampling error is the only factor contributing to *r*^2^.

## Methods

### Calculation of *r*^2^

For general use in analyzing LD with genotypic data, Weir (1979) recommended the composite (Burrows) disequilibrium statistic (Δ), which can be standardized to minimize effects of allele frequency, producing a correlation *r*_Δ_. Δ can be computed based on the distribution of multilocus genotypes, as described by Weir (1996). Then, *r*_Δ_ for loci 1 and 2 can be calculated as (Weir 1979, 1996):

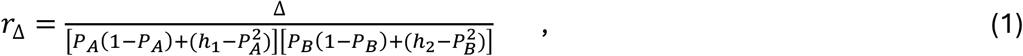

where *p*_*A*_, *p*_*B*_ are frequencies of alleles A (at locus 1) and B (at locus 2), *h*_1_ and *h*_2_ are observed frequencies of homozygotes for alleles A and B, respectively, and 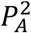 and 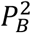are expected frequencies of those homozygotes. The terms 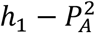and 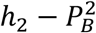 thus are adjustments for departure from single-locus Hardy-Weinberg genotypic proportions.

Subsequently, it was shown that if the genotypic vectors for two loci are coded 0/1/2 to reflect the number of copies of the focal allele every individual has, then the Pearson product-moment correlation of the two vectors is identical to Burrows’ *r*_Δ_ as calculated in Equation 1 (Gao et al. 2008). In what follows, we consider diallelic (∼SNP) loci, so there is a single 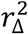value for each pair of loci.

### The expectation of 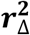

If selectively neutral alleles are tracked in a single closed population, the expected value of 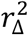 is a function of *N*_*e*_ and the recombination fraction (*c* = probability that two alleles will be shuffled during meiosis) (W&H):

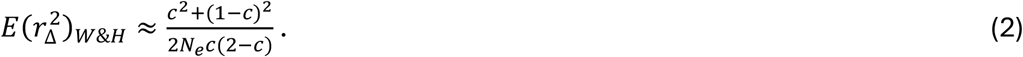

This equation accounts for contributions from drift and linkage (and their interaction) but not random error associated with sampling a finite number (*S*) of offspring. For the latter, W&H suggested that a term equal to 1/*S* could be added to Equation 2:

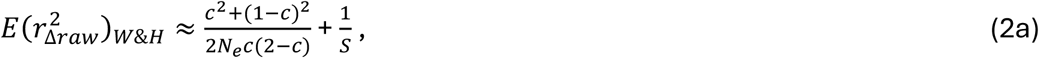

with the expectation now applying to LD in an empirical sample 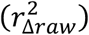rather than a population. It is noteworthy that W&H modeled the contribution of sampling error to 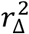 as strictly additive, independent of the variables *c* and *N*_*e*_:

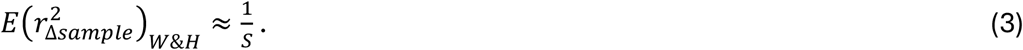

Consequences of this approach are discussed below.

The sample-size adjustment in Equation 3 was included by Hill (1981) in outlining the first formal approach for estimating *N*_*e*_ based on LD data. For any value of *c*, an estimate of the drift component to 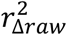 can be obtained as

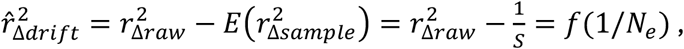

and the result can be rearranged to obtain an estimate of effective size. If analysis is restricted to pairs of loci on different chromosomes (so *c* = 0.5), effects of physical linkage are eliminated and Equation 2a simplifies to

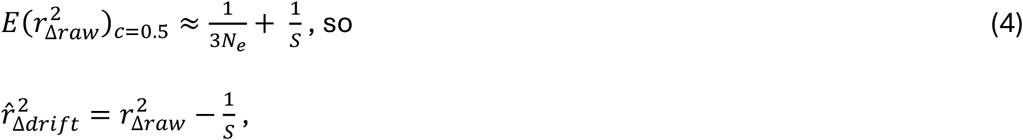

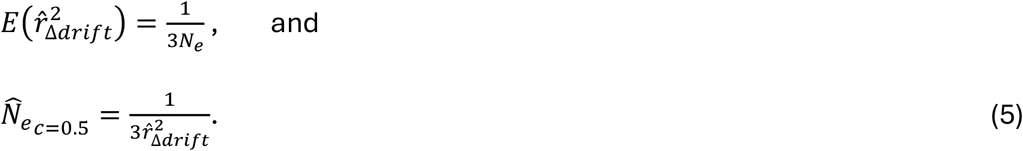

Equations 2-5 apply to unphased genotypic data (by far the kind most widely available for non-model species), and they are equally valid for species with separate sexes and monoecious species with random selfing (W&H).

Subsequently, it became clear that 1) Hill’s method can substantially underestimate contemporary *N*_*e*_ if sample size is smaller than effective size (England et al. 2006), and 2) this occurs because the actual contribution to 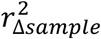 is larger than 1/*S* (Waples 2006). As a consequence of underestimating _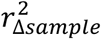,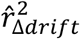_ is overestimated, leading to downward bias in 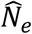.

To minimize sample-size related bias, Waples (2006) developed an empirical adjustment based on simulated data generated without any drift signal. First, he adopted Weir’s (1979) suggestion that an unbiased estimate of Δ can be obtained by multiplying the raw value by *S*/(*S*-1). This adjustment also applied to *r*_Δ_, so the raw 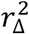 values Waples (2006) evaluated in developing his empirical bias correction were (compared to values based on Equation 1) higher by the factor [*S*/(*S*-1)]^2^. Next, because W&H ignored second-order terms in their derivations, Waples (2006) fit a function of 1/*S*^2^ to the residual 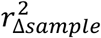 that remained after applying Hill’s 1/*S* sample-size adjustment to raw 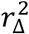. This resulted in expressions for 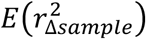 for two different ranges of sample size, and these adjustments were incorporated into the LDNe software (Waples and Do 2008) and eventually into NeEstimator2 (Do et al. 2014):

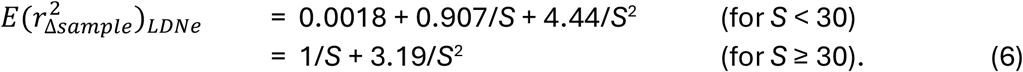

Although the empirical LDNe adjustments have been fairly effective in minimizing bias to 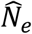 (Waples and Do 2010; Gilbert and Whitlock 2015; Wang 2016), they are not ideal for several reasons: 1) Since the W&H adjustment for sampling error (Equation 3) is too small even when *r*_Δ_ is calculated directly from Equation 1 (or using the *cor* function in R), it does not seem optimal to further increase empirical 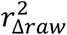 by multiplying it by [*S*/(*S*-1)]^2^. 2) Adding extra terms inversely proportional to *S*^2^ requires separate adjustments for small and larger samples. 3) The resulting quadratic terms can complicate *N*_*e*_ estimation.

Here I propose an alternative sample-size adjustment that, like Equation 4, is satisfyingly simple but, unlike Equation 4, accurately predicts 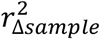 when no other factors contribute to LD:

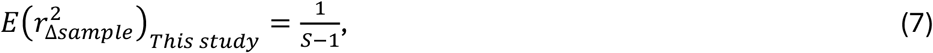

which is always >1/*S* for *S*>1. Using Fisher’s *k*-statistics, Pitman (1937) obtained the result in Equation 7 for the expected squared correlation between two random vectors of *S* elements, but that result seems to have been lost to history.

### Computer simulations

Numerical simulations (R core team 2026) were used to evaluate and compare accuracy of Equations 3, 6, and 7 for 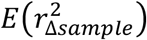. To isolate effects of sampling error, the simulations modeled an infinite effective size, which eliminated random LD due to drift. Although modeling large *N*_*e*_ is memory intensive and time consuming, infinite *N*_*e*_ can easily be modeled by drawing multilocus genotypes independently based on fixed allele frequencies. For each of 1000 ‘SNP’ loci, I drew the minor allele frequency randomly from the set [0.1, 0.2, 0.3, 0.4, 0.5]. Data were collected from sampled offspring numbering 10 to 500. Genotypes in samples were coded 0/1/2 as described above. For a locus with fixed minor-allele frequency *P*, offspring genotypes were 0 with probability (1-*P*)^2^, 1 with probability 2*P*(1-*P*), and 2 with probability *P*^2^. This mimicked reproduction in an infinite monoecious population. 50,000 replicates were evaluated for each sample size, and the empirical mean 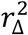 across locus pairs was compared with the three predictive formulae, as follows.

For each sample, genotypes were first filtered to remove monomorphic loci and loci for which every individual was a heterozygote (the correlation is not defined if one variable is invariant). Next, 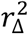 was calculated for all remaining locus pairs using the *cor* function in R. For the methods based on Equations 3 and 7, a mean residual 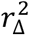was then calculated as 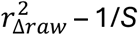and 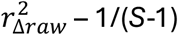, respectively. To mimic the approach used in the LDNe software, 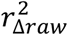 values were first multiplied by [*S*/(*S*-1)]^2^ (producing 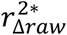 and then the residual 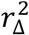 was calculated as 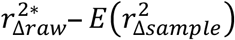, where 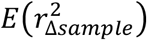 was based on Equation 6.

## Results

### Simulations

As expected based on previous results, the 1/*S* term proposed by W&H to account for sampling error is too small, resulting in a large residual 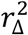 for small samples (top panel of Figure 1). On the scale required to display residual 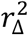 for the W&H method, which can exceed 0.01, residual 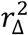 for the LDNe and 1/(*S*-1) methods are largely indistinguishable, both barely rising above the origin. Shrinking the scale of the vertical axis by over an order of magnitude (bottom panel of Figure 1) reveals that the LDNe method slightly overcorrects for sampling error (negative residual 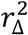) for *S* larger than about 40 and undercorrects (positive residual 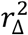) to a larger degree for the smallest samples. In contrast, the size adjustment based on Equation 7 (1/(*S*-1)) fully accounts for empirical 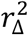 almost exactly, regardless of sample size.

**Figure 1.**
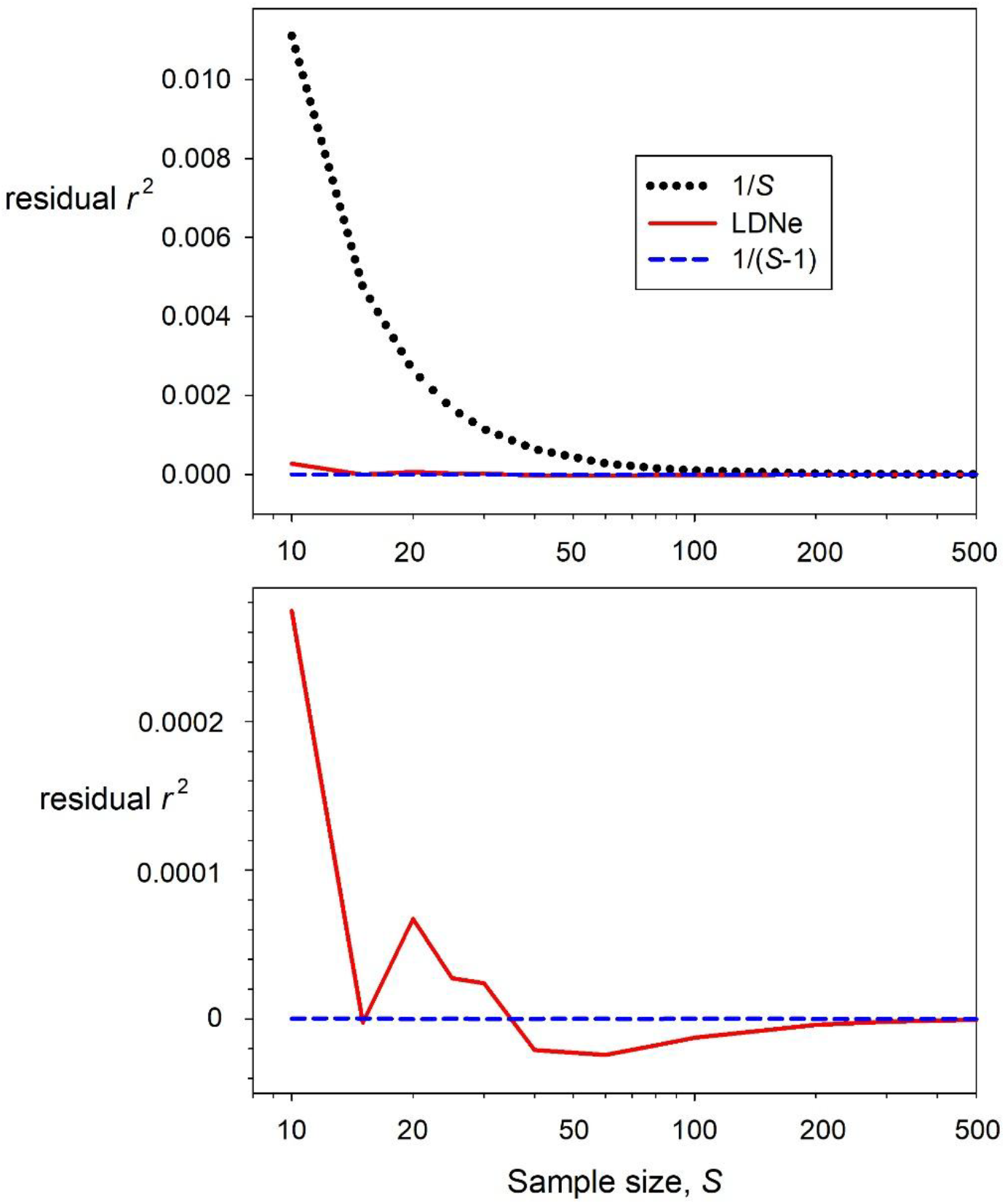
Results of simulations modeling contribution of sampling error to 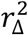 when *N*_*e*_ is infinitely large. Ideally, after accounting for sampling error, residual 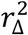 should be 0. Results are shown for three methods: 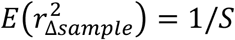 (W&H; Equation 3); 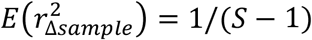 (this study, Equation 7); 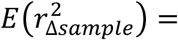 as in Equation 6 (LDNe). The bottom panel zooms in on the small departures for the two best methods. Note the log scale on the horizontal axis.

Figure 2 examines results for the 1/(*S*-1) method more closely, by rescaling the Y axis to values relevant for this method and by considering a range of allele-frequency distributions. For all 3 allele-frequency distributions, |residual 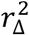 | values were tiny (<<10^-6^), the only exception being fixed parametric allele frequencies of 0.1, for which 10^-6^ < |residual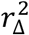 | <2×10^-6^ for the two smallest sample sizes (*S* = 10 and 15). Apart from this suggestion of higher variance for parametric *P* = 0.1 at the smallest sample sizes, residual 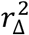was distributed approximately symmetrically around 0 (with both positive and negative deviations) for all allele frequency distributions, which is what one would expect if the 1/(*S*-1) term were independent of allele frequency.

**Figure 2.**
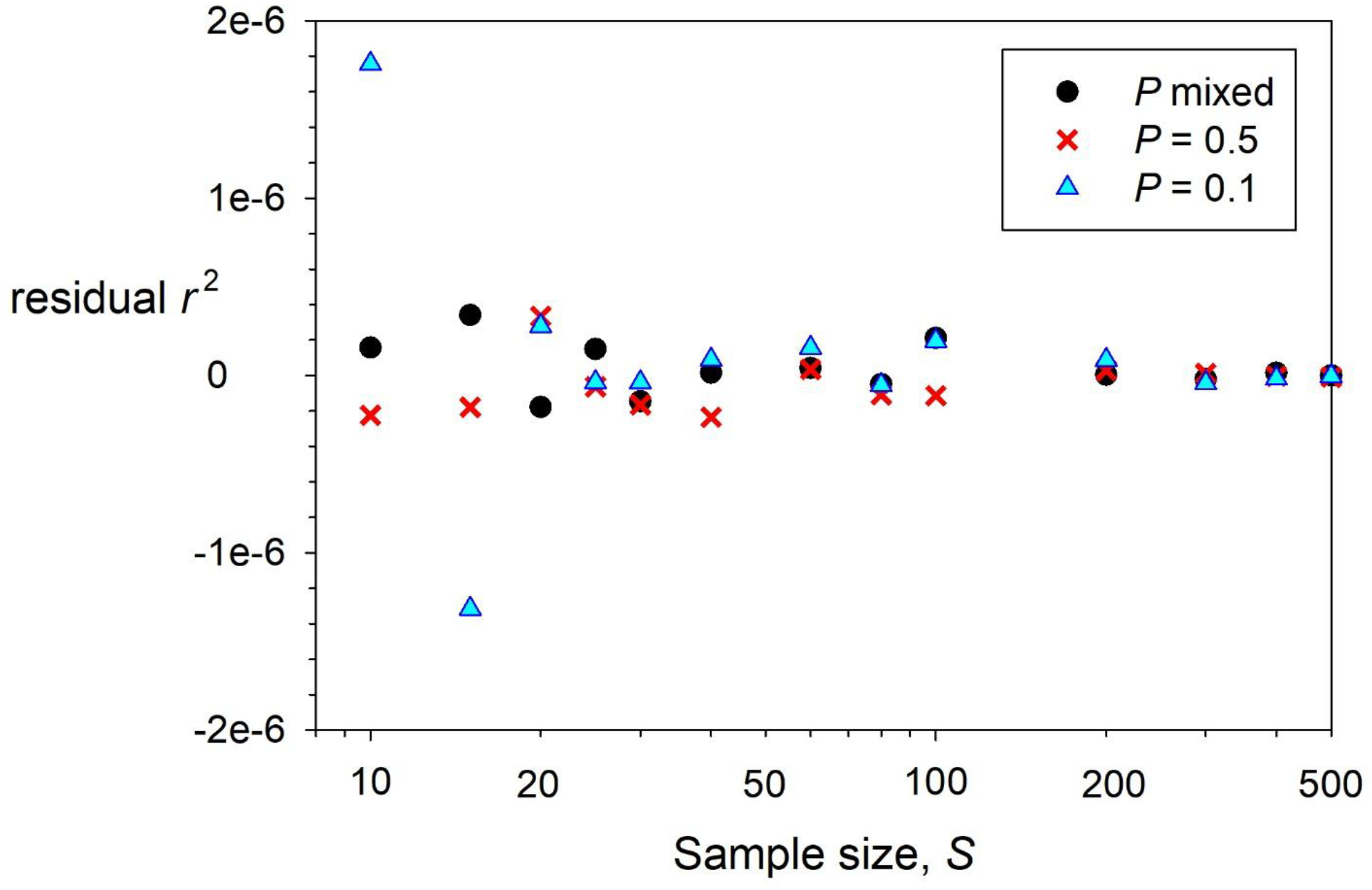
Detailed results for the 1/(*S*-1) method based on Equation 7, showing residual 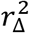 as a function of sample size, with the vertical axis scale focused on method-specific-relevant values. Colored symbols depict results based on different parametric allele frequencies. Note the log scale on the horizontal axis.

### Magnitude of sampling error when other factors contribute to 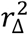

The above results are useful as a benchmark for evaluating performance in a scenario where effects of sampling error can be isolated from those due to additional factors, which include the four evolutionary forces (mutation, migration, selection, and genetic drift), as well as non-random mating and correlations due to physical linkage.

However, all real populations are finite, so even if the other factors are absent, some LD will be generated by genetic drift. Furthermore, many evaluations (e.g., GWAS studies for biomedical research) focus on locus pairs that are closely linked on the same chromosome and hence generate deterministic LD rather than the random LD generated by drift and sampling error. In general, therefore, analyses in real datasets will have to consider at least one additional component contributing to LD 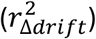 or two additional components (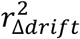 and 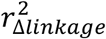), in addition to sampling error.

This is important because, in general, the absolute contribution of sampling error to empirical 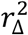 must shrink as background LD becomes stronger. At the extreme, perfect LD within a population 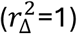 occurs when an individual’s genotype at one locus of a pair reliably predicts its genotype at the second locus. In that scenario, any possible subsample of individuals also produces 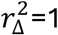, so 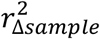 shrinks to 0. This effect is apparent in Figure 1 of Bercovich et al. (2026), which shows the decline of what is referred to here as 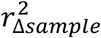 as the underlying correlation between two loci increases.

This is a complicated topic that merits more detailed evaluation than can be attempted here. However, we can gain some insight from another published result that bears on issues considered here. In the Appendix to Bulik-Sullivan et al. (2015b), the following equation appears without explanation or attribution:

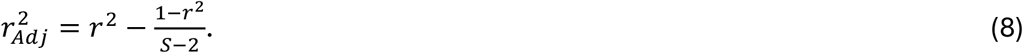

A possible source for this equation is Ezekiel (1930), where a related formula appears on p. 267. Equation 8 can be recast in current notation as

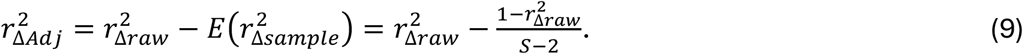

In this formulation, 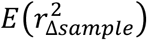 is a decreasing function of 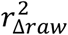 and shrinks toward 0 as 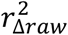 approaches 1.

We can evaluate the relationship between Equation 8 and Equation 7 when sampling error is the only contribution to 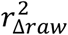. In that case,

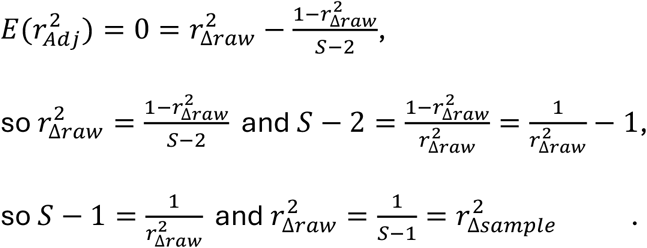

Thus, the Bulik-Sullivan (2015b) equation correctly gives 1/(*S*-1) as the contribution to 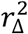 from sampling error when all other contributions are absent.

## Discussion

As noted by Bercovich et al. (2026), it probably is not feasible to expect a completely unbiased general expectation for *r*^2^. Based on results presented here, however, it is possible to exactly specify *E*(*r*^2^) for an important special case, where the only contribution to *r*^2^ is from sampling a finite number of individuals.

The relative importance of this result will vary depending on the research goals. Biomedical researchers searching for associations between molecular loci and diseases or specific phenotypes, and scientists interested in historical demography, both commonly focus on tightly linked genes that are deterministically expected to produce relatively large *r*^2^ values. In those scenarios, the actual contribution of sampling error could be substantially less than the simple expression in Equation 7, thus necessitating some sort of adjustment akin to what is done in Equation 8.

At the other extreme, the LD method to estimate contemporary *N*^*e*^ using pairs of unlinked loci is only concerned with 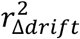 and 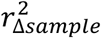, and even when both are relatively small, empirical 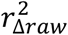 will generally also be small (<0.1). For example, with *S* = 20 and *N*_*e*_ = 50 (lower limit of the ‘50-500’ rule for population viability), 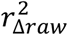 would be expected to be about 1/(*S*-1) + 1/(3*N*_*e*_) = 0.0527 + 0.0067 = 0.0593. When 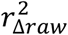 is this small, there is very little shrinkage of the sampling error component (Figure 3), so Equation 7 should be 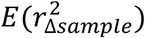 effective as a general estimator of *E*(*r*^2^).

**Figure 3.**
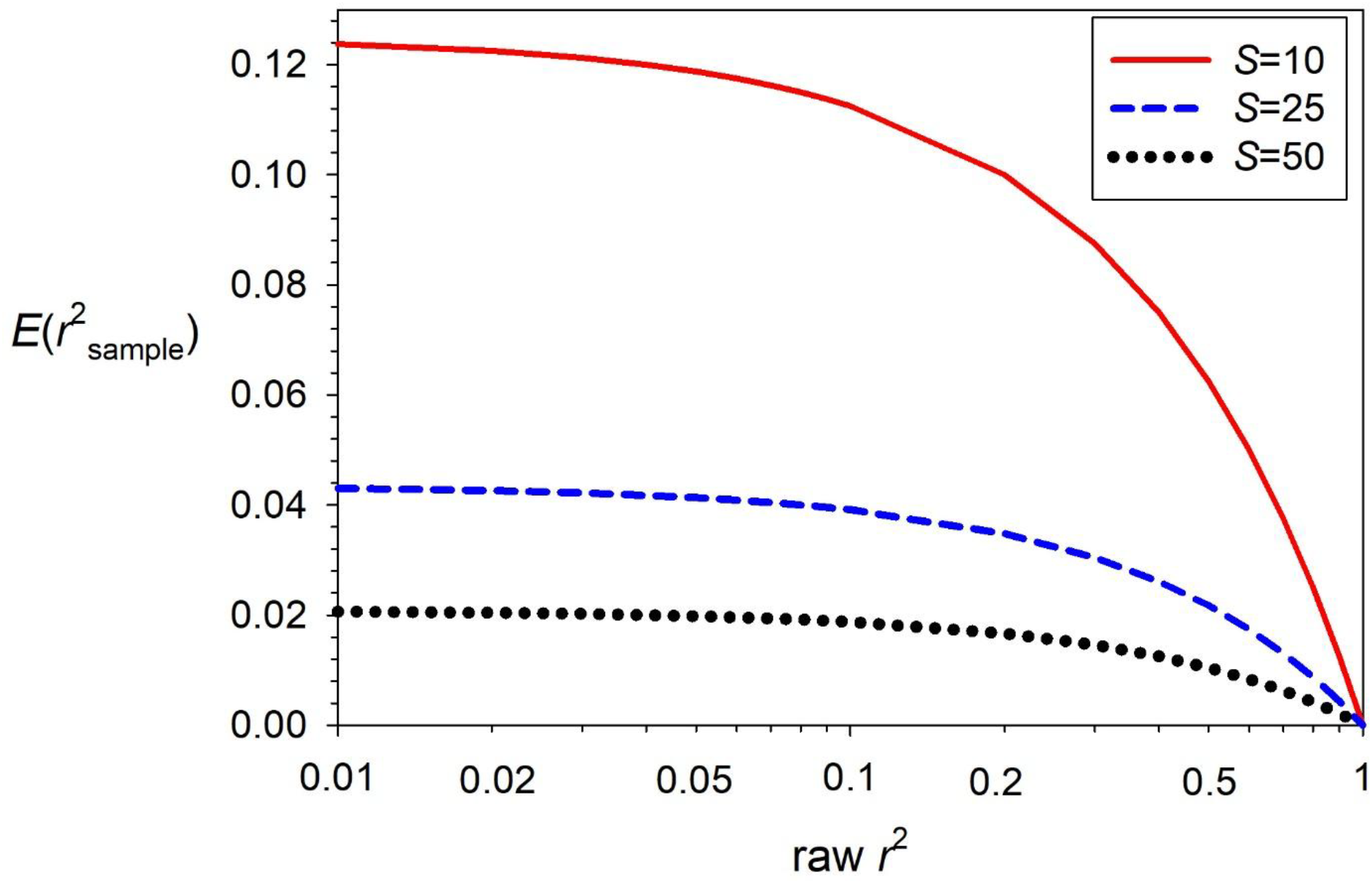
Theoretical shrinkage of 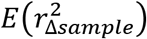 as empirical 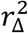 increases, for three different sample sizes. 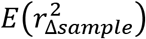 was computed as the rightmost term in Equation 8, from Bulik-Sullivan et al. 2015b. Note the log scale on the horizontal axis.

## Acknowledgments

The original motivation for exploring Equation 7 came from Sved et al. (2013), who presented a number of equations with terms in 2*S*-1 to account for sampling error. This manuscript benefitted from discussions with Anders Albrechtsen and Ryan Waples, and I am grateful to them for archeological work uncovering the origins of obscure equations.

---

**Table 1.** Notation used in this study.

|  |  |
| --- | --- |
| $S$ | Number of individuals sampled |
| $N_e$ | Effective population size |
| $P$ | Allele frequency |
| $h_1, h_2$ | Frequency of homozygotes at locus 1 and locus 2 |
| $c$ | Recombination fraction ($0 \leq c \leq 0.5$) |
| W&H | Weir and Hill 1980 |
| $\wedge$ | Denotes an estimate |
| $E$ | Denotes an expectation |
| $\Delta$ | Burrows' version of the disequilibrium coefficient $D$ |
| $r_\Delta$ | A form of $\Delta$ standardized by allele frequency to produce a correlation |
| $r_\Delta^2$ | The square of $r_\Delta$ = squared correlation of alleles at two loci |
| $r_{\Delta raw}^2$ | Empirical (observed) value of $r_\Delta^2$ |
| $r_{\Delta raw}^{2*}$ | $r_{\Delta raw}^2$ adjusted with the $[S/(S-1)]^2$ factor suggested by Weir (1979) |
| $r_{\Delta sample}^2$ | Contribution of sampling error to $r_{\Delta raw}^2$ |
| $r_{linkage}^2$ | Contribution of reduced recombination ($c < 0.5$) to $r_{\Delta raw}^2$ |
| $r_{\Delta drift}^2$ | Contribution of genetic drift to $r_{\Delta raw}^2$ |
| $\hat{r}_{\Delta drift}^2$ | An estimate of $r_{\Delta drift}^2$; $\hat{r}_{\Delta drift}^2 = r_{\Delta raw}^2 - E(r_{\Delta sample}^2)$, assuming other factors contributing to $r_{\Delta raw}^2$ are absent |
| residual $r_\Delta^2$ | Difference between $r_{\Delta raw}^2$ and $E(r_{\Delta sample}^2)$ |
| $r_{Adj}^2$ | In Bulik-Sullivan et al. (2015b), $r_{\Delta raw}^2$ after accounting for expected contribution of sampling error |

